# Tripal Developer Toolkit

**DOI:** 10.1101/328641

**Authors:** Bradford Condon, Abdullah Almsaeed, Ming Chen, Joe West, Margaret Staton

**Affiliations:** University of Tennessee Knoxville, Entomology and Plant Pathology; University of Tennessee Knoxville, Genome Science and Technology

## Abstract

Tripal is an open-source biological community database construction toolkit utilizing the content management system Drupal. Tripal is used to make biological, genetic and genomic data more discoverable, shareable, searchable, and standardized. As funding for community level genomics databases declines, Tripal’s open source codebase provides a means for sites to be built and maintained with a minimal investment in staff and new development. Tripal is ultimately as strong as the community of sites and developers that use it. We present a set of developer tools that will make building and maintaining Tripal 3 sites easier for new and returning users. These tools break down barriers to entry such as setting up developer and testing environments, acquiring and loading test datasets, working with controlled vocabulary terms, and writing new Drupal classes.

## Introduction

The rapid advancement in sequencing technology has resulted in a proliferation of genomic data. General, all-purpose databases such as NCBI capture some of this data, but additional support is needed for manual annotation, specialized analyses, and data integration, particularly for groups not specializing in bioinformatics. Community-level genomics databases fill this role, hosting a curated set of data from a species or multiple species of interest.

In recent years, however, funding for these resources has been significantly reduced, even for large-scale model organism databases (1). To continue the critical role they play connecting researchers to tailored bioinformatic resources, community databases must formulate a plan to not only keep sites available but also continue to add new data and services. Additionally, data repositories are now striving to meet new accessibility and metadata standards for public data that span four categories: findable, accessible, interoperable and reusable (FAIR) (2,3).

The biological community database construction toolkit Tripal was created as a multifaceted solution to many of these problems (4,5). Tripal marries the content management system Drupal (http://www.drupal.org) to the standard biological relational database storage backend Chado (6). Tripal utilizes the same system of modular code units that have made Drupal so successful. Tripal core provides a basic set of common functionality, including content types such as organisms, sequence features, and controlled vocabularies. Extension modules can then be developed by any group to extend the core and provide additional functionality. Examples of extension modules include a BLAST tool for users (7), natural diversity genotype data loader and display (8)(9), and elasticsearch to provide fast sitewide searching (10). Tripal websites cost less to build and maintain, and they can reuse and share Tripal extension modules, meaning each website is not coding its own solution to shared biological problems. These advantages make Tripal an attractive option for community database platforms aiming to do more with fewer resources while remaining sustainable.

Tripal was first released in 2009 and has since had numerous improvements (4,5), the latest of which, Tripal 3 (11), includes improvements to interoperability and FAIR accessibility. For example, Tripal sites now link all data to Controlled Vocabulary (CV) terms. Tripal 3 also upgrades many of the Drupal concepts, a necessary task as Drupal’s release cycle marches forward. In particular, nodes are replaced with **Bundles, Entities**, and **Fields** to allow more lightweight, flexible content type solutions (Table 1).

**Table 1.** Key Drupal vocabulary and concepts. An entity type is essentially a generic container that defines a type of content, while a bundle is a more specific subtype that includes fields. For example, Drupal has a page entity type, with bundles for more specific types of pages, for example a blog post and an article. These bundles include fields that define a smaller unit of content associated with a bundle. In the case of the blog post, fields would include a title, the author, the date it was posted, and the body of the article. A entity is a piece of specific content, such as a specific blog post. In Tripal, examples of bundles would include units of content such as gene or organism. Fields would include gene name or genus and species. A single species, such as *Fraxinus excelsior,* would a be an entity.

| Drupal Concept Introduced in Tripal 3 (see http://tripal.info/node/347) | Definition | Example | Corresponding Tripal 2 concept |
| --- | --- | --- | --- |
| Entity Types (Tripal Entity Type) | A group of fields. All Tripal content is of the Tripal Entity type | Tripal Entity Type, Page | Node |
| Bundle (Tripal Content Type) | An implementation of an entity type to which fields can be attached. | Organism | Node Content Type |
| Field | A piece of content that can be used in multiple bundles. | Common name, Genus, Species |  |
| Entity | A particular instance of a Tripal Content Type. | The entry *Fraxinus excelsior* | Node |

### Developer Challenges

The Drupal website lists 121 sites using Tripal as of this writing (https://www.drupal.org/project/usage/tripal), a number which continues to grow. As more communities rally behind Tripal, we expect the developer community to grow alongside it. While it is advantageous that Tripal sites can be created with a smaller staff, this also introduces its own set of challenges. Creating custom Tripal modules often requires some understanding of biology, as well as significant expertise in computer science and web development. Maintaining the site (collecting, curating, and uploading data) requires skills in web development, biocuration and bioinformatics. Development teams may not have the personnel to fulfill all these roles.

Tools that lighten the workload and ease the learning curve should be very welcome for new and veteran Tripal module developers alike. We recognize that the skillset of a given development team may be lacking in biological or computer science expertise. Given that Tripal lives at the union of these disciplines, we identify challenges a small development team is likely to face deploying a Tripal site, and present a suite of developer tools to facilitate their resolution (Table 2) which we have developed in the course of our own work on the Tripal-based site Hardwood Genomics Project (12).

**Table 2.**
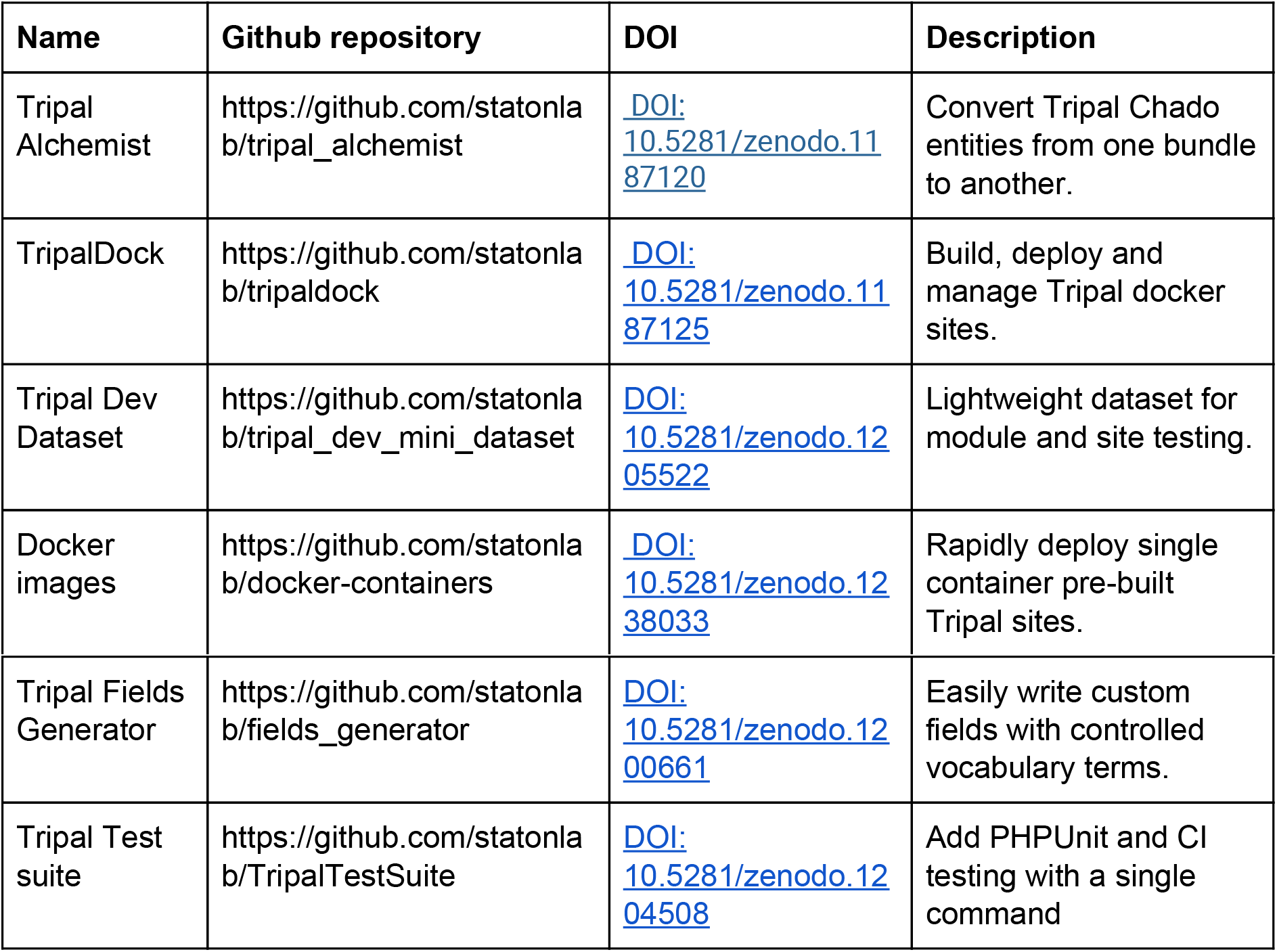
Tools presented in this paper.

#### Developer instance

Developers must set up a developer instance to test code before deployment. Building, maintaining, and rebuilding developer sites is time consuming and requires system administrator knowledge that may be beyond developers with primarily biology backgrounds. Docker containers help developers quickly and easily create shareable development environments, and using them is developer best practice (13). **TripalDock** automates and optimizes these tasks.

#### Data acquisition

Development sites should be lightweight, but they still require biological data to test loading, display, and manipulation of real data. Computer-science focused developers may have difficulty understanding how to generate, format, and load Chado with complicated interconnected bioinformatics datasets. **Tripal DevSeed** is a curated, miniature dataset in one place, covering all core Tripal functionality.

#### Testing and Continuous Integration

Code testing is widely appreciated by professional software developers for the many benefits it provides (14). Tests ensure that software produces the intended result given various inputs, and their use can help discover errors or bugs in the code. Tests also ensure that previously written code is not broken by new additions, and they allow safe code refactoring. The most common testing framework for PHP, the coding language of Drupal and Tripal, is PHPUnit (15). As with most testing frameworks, it uses “assertions” to verify that code behaves as expected. Consider a method that renames a gene feature. A suite of tests might verify that given input features are correctly renamed, and that the method can gracefully deal with problematic cases such as when the gene does not exist in the database, or the new name is already in use. If the rename method is updated with new code, tests are rerun to ensure no new bugs have been in inadvertently created by the new additions. Tests are so helpful that some development practices prescribe writing tests before writing the intended code (Test Driven Development), which can increase test coverage and coding efficiency (10). Despite their importance, a novice Tripal developer may be intimidated by the overhead required for writing reproducible, platform independent tests that interact with Chado, Drupal, and Tripal.

Continuous integration (CI) is the practice of frequently merging new code during development, then automatically building and testing the new codebase to verify integrity software. Adding CI testing to software allows projects to release updates more often (16), and to included more code from non-core developers without diminishing code quality (17). One good solution for Tripal developers to leverage is Travis CI, a free service to build and test code hosted at GitHub (18). Testing and CI is a huge topic, and biology-focused developers may not be familiar with it or why it is important to learn and use. The Tripal Test Suite encourages developers to learn about these software development practices and eases their initial set up by automatically adding the framework for PHPUnit and Travis CI to a Tripal module.

#### Entity management

Tripal code developers need to contend with new Drupal content type concepts, including bundles, entities, and fields. Tripal ships with a migration function to convert core node types (an earlier Drupal content type) to entities, but there is no tool to convert entities from one type to another. Additionally, in Tripal 2, many extensions provided custom node types. For example, analysis subtype modules defined BLAST and Interproscan analyses. The Tripal 3 migration process cannot support all custom nodes, so they migrate as analysis entities, and must be converted to custom entities to preserve unique functionality. As site administrators build their sites, they will likely want to further customize content types. **Tripal Alchemist** is an easy to use tool for this task.

#### Coding fields

Fields are an integral component of the new entity system. In simple cases, new fields can be easily created through the Drupal administrative web pages. However, defining fields programmatically requires an understanding of Chado, Controlled Vocabularies, and the Tripal field structure. Furthermore, multiple, interconnected files with strictly defined function naming are required, leading to simple mistakes that are difficult to troubleshoot. The **Tripal Fields Generator** automates the field creation process.

The tools we have developed to facilitate our own Tripal development are open source and freely available, and we hope they will accelerate the development and sharing of new Tripal modules for others as well

## Materials and Methods

### Code standards and accessibility

All projects and tools are available on github under an open source license (GPLv3 or MIT). Each individual project includes documentation for installation and use, as well as guidelines for contribution.

### Tripal DevSeed Sequences and Annotations

Two hundred CDS sequences and their corresponding predicted amino acid sequences of the *Fraxinus excelsior* genome assembly were used to seed the developer dataset (19). CDS sequences were annotated using BLAST (20) against UniProtKB-TrEMBL and the plant portion of UniProtKB-SwissProt (21). Amino acid sequences were annotated using InterproScan 5.4 (22) using the iprlookup, goterms, and pathways flags. Biosamples were created by randomly modifying existing *Fraxinus excelsior* NCBI Biosamples (23). Expression data was randomly generated for each Biomaterial and CDS pair. The Newick format tree file was generated by aligning CDS sequences using MAFFT (24). Kyoto Encyclopedia of Genes and Genomes (KEGG) annotations were generated using the online KEGG Orthology And Links Annotation (KOALA) tool (25).

## Results

### Developer instance tools

The first step in developing any sort of web application is setting up a developer environment. A development environment preserves the functionality of your live site for your users, protects your data from accidental corruption, and, if configured well, can greatly speed up your development time. For a Tripal site, a developer instance must consist of a **webserver** (apache/nginx) and **database** (PostgreSQL) with Tripal’s base dependencies installed (PHP, Drupal, PHP extensions) and configured correctly, as well as any additional services (phppgadmin or elasticsearch). While the Tripal user’s guide provides manual installation instructions (available at http://tripal.info/tutorials/v3.x/installation), this can be an overwhelming experience for new Tripal developers without prior web development experience.

We present two platforms for rapid developer instance deployment. The first is a self contained docker image, with webserver and database. Docker provides a way to “containerize” a software program or set of software programs, allowing them to be quickly and easily transferred among computing systems and re-deployed without any new configuration or installation effort. We have created docker images for base Drupal (no Tripal installed), Tripal 2, and Tripal 3. The Tripal 3 image is also provided pre-loaded with a miniature dataset stored in Chado (described below). For those experienced with docker, the provided images allows single-command deployment of a functional Tripal site. Because they are already initialized, deployment is extremely fast, taking seconds rather than minutes. For developers that need to build more customized and permanent containers and/or without prior knowledge of Docker, we provide a second solution, TripalDock. As opposed to the Tripal images already built and described above, TripalDock is a command line tool that enables developers to create and interact with their own custom Docker containers. It provides simple commands to perform cumbersome Docker tasks, and handles the mapping of files, creation of containers, execution of admin commands, dependency management, and volume cleanup (Table 3). This tool allows new developers to get started immediately, avoiding the pitfalls of setup and configuration that can stump novice programmers.

**Table 3:** Commands provided by Tripaldock.

| Command | Description |
| --- | --- |
| new | Create a new container. |
| up | Start the Tripaldock container. |
| down | Stop the Tripaldock container. |
| rm | Remove and destroy the Tripaldock container. |
| ssh | Interactive shell access to the container. |
| logs | Access the logs of various services running inside the container. |
| install | Equivalent to `drush pm-enable tripal_module`. Allows users to install modules from outside the container. |
| drush | Run drush commands from outside the container. |

### Developer Dataset

Community genomics database house and manipulate large sets of interconnected biological data. Developers, particularly those who lack a background in biology and bioinformatics, can quickly become frustrated locating, formatting, and importing the appropriate biological data into their developer site to develop and test their code. Mirroring the dataset available on their live website may be impossible due to the size of genomic datasets, which can result in extremely large databases that take up too much storage space and run too slowly for the rigors of development.

We therefore generated a truncated dataset of 200 genes from the genome assembly of F. *excelsior* (19). Data types appropriate for all of the core and many common extension Tripal modules are available (Table 4) as a set, called Tripal DevSeed. Docker images can be downloaded with the data preloaded, or, developers can opt to download clean images and upload only the data relevant for their work. Detailed guides for loading data are included in the GitHub repository.

**Table 4.** Developer Dataset contents

| Name | Corresponding module | Format | Description |
| --- | --- | --- | --- |
| sequences | Core | FASTA | 30 Scaffolds with 200 CDS sequences and their corresponding polypeptide sequences. |
| gff | Core | GFF | GFF3 describing gene, mRNA, exon, CDS, and five/three prime UTR regions of the above 30 scaffolds. |
| blast_annotations | tripal_analysis_blast | XML | BLAST annotations for CDS sequences against swissprot and TREMBL databases |
| interposcan_annotations | tripal_analysis_interpro | XML | Interproscan annotations of amino acids. |
| kegg_annotations | tripal_analysis_kegg | TSV | KEGG annotations generated by the kegg koala tool. |
| biosamples | tripal_analysis_expression | ncbi xml | Two sets of 3 and 4 NCBI XML formatted biosamples. |
| expression_data | tripal_analysis_expression | TSV | Randomly generated expression data for biosample set 1 and 2. |
| database_backups | NA | SQL | postgresql database dump of all loaded biomaterials, facilitating one-step loading of a database. |
| fexcelsior_mini_set.tree | Core | Newick | Newick format tree generated from MAFFT-aligned CDS |

### Testing and Continuous Integration

Testing is an essential step in software development and particularly important for shared community code development, yet many projects do not adopt a formal testing structure, typically because developers are pressured to provide results and feel they cannot afford the time spent learning a framework and writing tests. **Tripal Test Suite** lowers the barrier to entry for writing PHPUnit tests for Tripal: installing the package allows single command set up of the necessary directory structure, environmental variables, and database connection (see Table 5 for a full list of features). Tests can be wrapped in a database transaction and rolled back automatically, preserving your database. **Data factories** allow users to easily add biological data into Chado that can be altered in the test and rolled-back after it finishes.

**Table 5.**
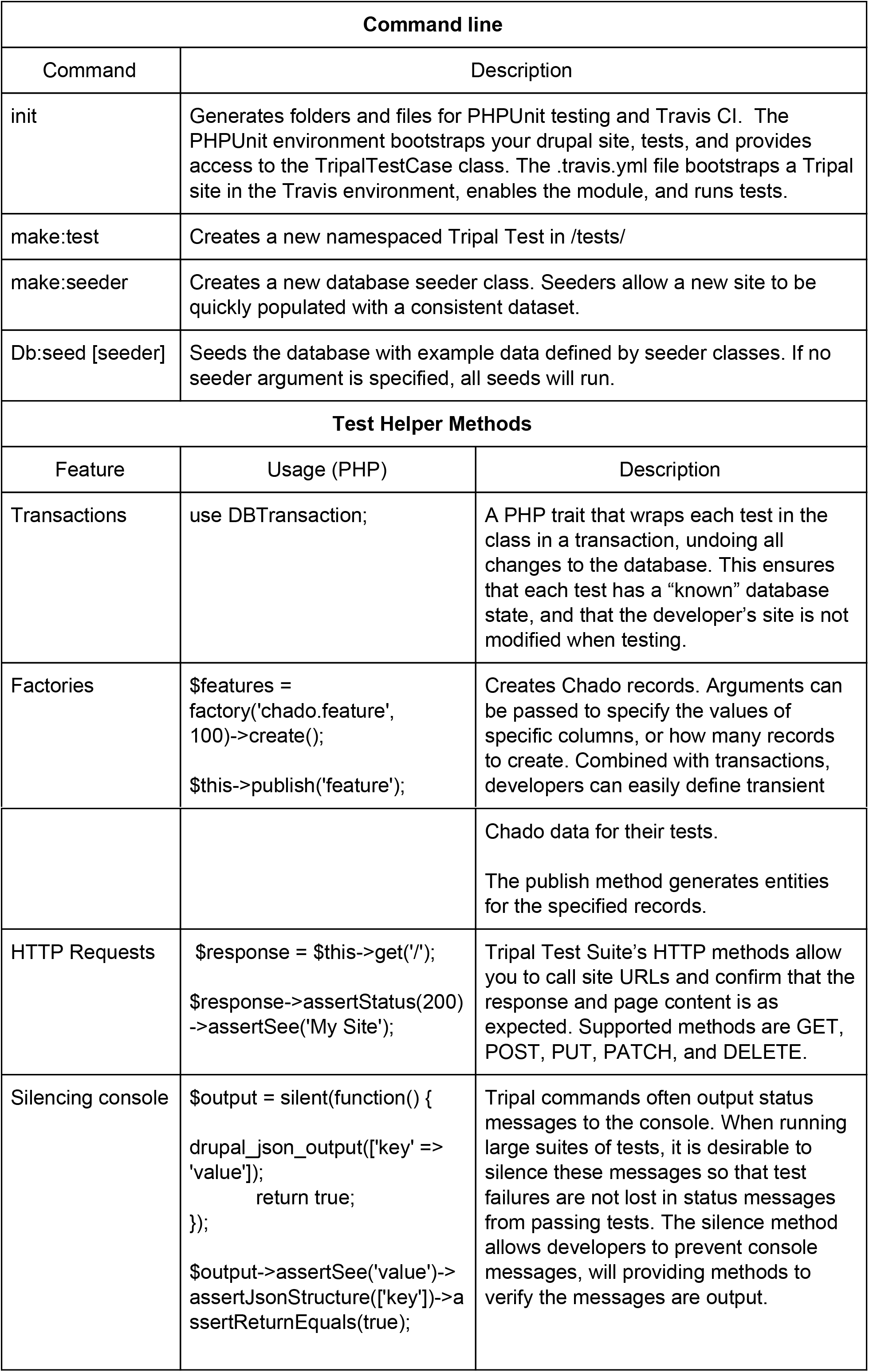
Tripal Test Suite Features.

| Command line |  |  |
| --- | --- | --- |
| Command | Description |  |
| init | Generates folders and files for PHPUnit testing and Travis CI. The PHPUnit environment bootstraps your drupal site, tests, and provides access to the TripalTestCase class. The .travis.yml file bootstraps a Tripal site in the Travis environment, enables the module, and runs tests. |  |
| make:test | Creates a new namespaced Tripal Test in /tests/ |  |
| make:seeder | Creates a new database seeder class. Seeders allow a new site to be quickly populated with a consistent dataset. |  |
| Db:seed [seeder] | Seeds the database with example data defined by seeder classes. If no seeder argument is specified, all seeds will run. |  |
| Test Helper Methods |  |  |
| Feature | Usage (PHP) | Description |
| Transactions | use DBTransaction; | A PHP trait that wraps each test in the class in a transaction, undoing all changes to the database. This ensures that each test has a “known” database state, and that the developer’s site is not modified when testing. |
| Factories | `$features = factory('chado.feature', 100)->create(); $this->publish('feature');` | Creates Chado records. Arguments can be passed to specify the values of specific columns, or how many records to create. Combined with transactions, developers can easily define transient |
|  |  | Chado data for their tests.<br>The publish method generates entities for the specified records. |
| HTTP Requests | `$response = $this->get('/'); $response->assertStatus(200) ->assertSee('My Site');` | Tripal Test Suite's HTTP methods allow you to call site URLs and confirm that the response and page content is as expected. Supported methods are GET, POST, PUT, PATCH, and DELETE. |
| Silencing console | `$output = silent(function() { drupal_json_output(['key' => 'value']); return true; }); $output->assertSee('value')-> assertJsonStructure(['key'])-> assertReturnEquals(true);` | Tripal commands often output status messages to the console. When running large suites of tests, it is desirable to silence these messages so that test failures are not lost in status messages from passing tests. The silence method allows developers to prevent console messages, will providing methods to verify the messages are output. |

The package also automates the configuration of **Continuous Integration (CI)** with Travis CI: the user need only push their code to GitHub and integrate their repository on Travis-CI. Out of the box CI will ensure the module can be installed on a Tripal 3 site and that all PHPUnit tests pass. Tripal Test Suite also provides support to automate the process of inserting and removing test data into the database using database seeders, which make test data available only during the testing process (Note that “auto increment” columns are not reset to their previous state). This concept is useful because it allows tests to manipulate the database without causing side effects by automatically rolling back the database state upon the completion of tests or if a fatal error occurs during the testing runtime. By secluding data providers from tests, database seeders also make it possible to share code that populates the database with mock data with other modules that depend on the same data.

The Tripal Test Suite is now used within the Tripal Core repository itself. Using the same tool for modules and core makes it easier to write and structure tests.

### Entities, Bundles, and Fields

Tripal 3 introduces the concepts of entities, bundles and fields. The resulting content is more flexible, and, because bundles and fields are associated with CV terms, more FAIR. This is another concept to master to successful develop in Tripal 3 and choosing CV terms can be difficult, especially for non-biologists. We present **Tripal Alchemist** and **Tripal Fields Generator** as essential tools to help developers work with bundles/entities and fields, respectively.

#### Tripal Alchemist

**Tripal Alchemist** is an administrative tool that facilitates converting Tripal entities representing Chado records from one bundle type to another (Figure 1). The module provides three avenues for converting entities. *Automatic* conversion will identify records whose type matches a different bundle. Analysis records, for example, migrate during the Tripal 2 to Tripal 3 upgrade process as a generic analysis bundle type, but they may have a specific type property. When the site administrator creates an appropriate bundle (BLAST annotation analysis type for example), matching entities will automatically be converted. *Manual* conversion provides a table for users to select entities to convert, overwriting their type. This is useful when creating a new bundle type that applies to existing records which do not have the matching property. *Collection* conversion functions similarly to *manual,* but acts on a collection of entities that the administrator can define elsewhere. This method is ideal for converting large subset of entities. For example, we wished to convert the mRNA records for some (but not all) species to mRNA contigs, allowing us to customize the fields displayed for these records.

**Figure 1.**
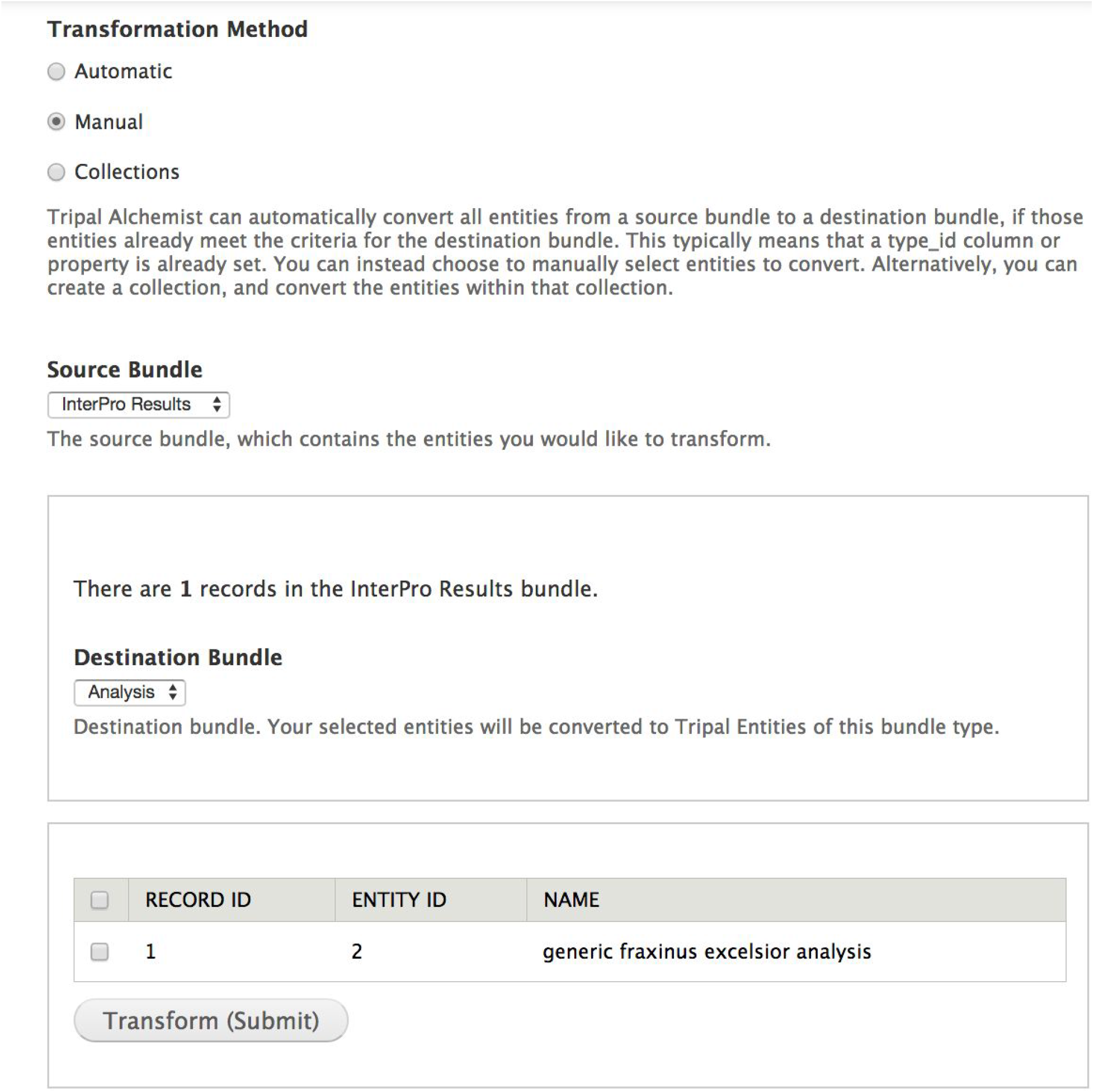
The Tripal Alchemist administrator tool. The Tripal Alchemist administrator interface. To convert entities, the administrator must first select a transformation method (top). In this example, choosing the manual method allows conversion of selected entities from a source type to a destination type (middle). The available source entities are then presented in a table (bottom) which the user can select from and submit.

Any information a developer wants displayed on the website must correspond to a field. A properly coded field should handle querying and retrieving the data from the storage backend (Chado, if a Tripal Chado field), providing an interface for the user to enter data into the field when creating or ending the bundle it is attached to, as well as formatting the data for both web services and end user consumption. Furthermore, fields are CV-centric: each field must correspond uniquely to a CVterm.

#### Tripal Fields Generator

We created the **Tripal Fields Generator** tool (TFG) to make coding fields significantly faster and easier. TFG is a command-line PHP package easily installed with composer. It asks a series of questions about the CVterm usage of your field and generates stub files with base class methods and instructions for both TripalFields and ChadoFields (Figure 2). Three files are generated to properly define the field, with standard functions already named and left empty for the developer to add code: the class file, the formatter file, and the widget file. Additionally, a stub file is generated demonstrating how to declare instances of the new field within the module. Because the cross-file references are generated automatically, this significantly reduces difficult to debug errors. Furthermore, TFG will connect to your Drupal site’s database automatically and verify that the CVterm exists in your database.

**Figure 2.**
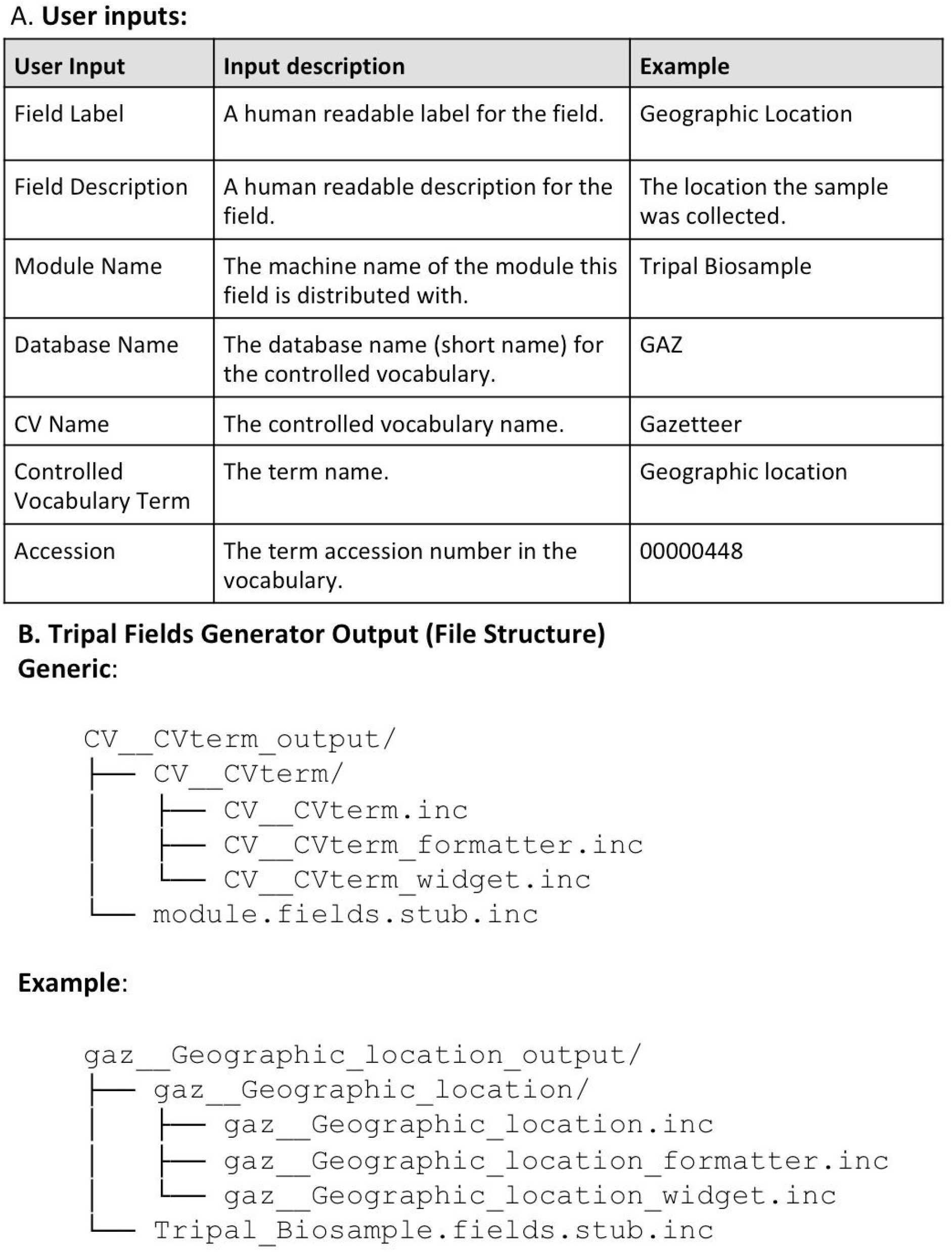
Input and output for Tripal Fields Generator. The command line program prompts the user to provide a series of inputs (A) and checks the database to determine if the specified controlled vocabulary term exists. If it does, the software produces a set of files in the correct directory structure to generate the field (B). An example is given using the “Geographic location” term from the Gazetteer vocabulary for a mock module named Tripal_Biosample (C). This demonstrates how the user’s inputs are structured into the file names. These inputs are also used to build all the necessary code variables and functions in each file, leaving only the essential custom coding for the function bodies for the developer to finish.

### Use Cases

To illustrate how a Tripal developer might utilize these tools in their own work, we have developed a set of three use cases, each of which highlights how one or more of the dev tools can be used.

#### Upgrading from Tripal v2 to Tripal v3

A developer is tasked with upgrading an existing Tripal 2 site to Tripal 3. After performing the built-in migration, all analyses are now using the same generic analysis bundle, and all analyses are displayed the same way. The developer wishes to customize certain analyses, in particular, the BLAST and InterProScan analyses that have been run for sets of genes. When the developer defines new bundles for these two types of analyses, they can use **Tripal Alchemist** to automatically convert the generic entities to the new, custom bundle types based on their matching type CVterms.

The developer wants to bring their Tripal 2 custom functionality to Tripal 3. They are working with a custom module called Tripal Sample, and they want to include a custom pane that displays the geographic location of a sample on a map. They use **Tripal Fields Generator** to define a new field, picking a CVterm for their field that exists in an ontology on their site. They pick the EDAM term “Geographic Location”, but forget that the accession is data:3720, using EDAM:3720 instead. Without Fields Generator, they would have struggled getting their field to work, or inserted a bad term into their site. However, Tripal Fields Generator checked the database and informed them of their mistake before it was made. The tool creates the three field stub files for them (data__geographic_location.inc, data__geographic_location.formatter.inc, data__geographic_location_widget.inc) and the code to declare the new field (tripal_sample_module.fields.inc), greatly reducing the possibility of making a very difficult to troubleshoot mistake (Figure 2).

#### Developing a new module

A developer is tasked with adding new functionality to their site: a genetic cross viewer to display F1 and F2 progeny phenotypes and genotypes. The first thing they might do is use **TripalDock** to create a new development site, so that they can work locally and not accidentally bring down their development site as they work. The developer is interested in displaying their cross data in the context of other data: phenotypes might link to genes and expression data. They therefore use **DevSeed** to quickly load in an organism complete with a miniature genome, biomaterials, and expression data.

The developer begins in earnest on their module. They use Tripal **Test Suite**’s init method to automatically set up a PHPUnit testing environment and continuous integration. As the developer starts work on an importer for cross data, they write unit tests for functionality in the importer. The developer might work with a biologist at this stage to define the expected constraints on the data, such as: “Can an individual appear in the same cross multiple times?”, “Do all progeny always have both a phenotype and a genotype?”. These constraints can be included in tests to ensure the biologist and the developer both achieve consistent and expected functionality. Each test can be wrapped in a database transaction, so new methods being tested don’t alter the database for older, working tests. When the importer is completed, the developer can easily refactor their code, simply re-running the unit tests after each refactor to ensure functionality.

The developer starts coding a field to display the cross data. They use **Tripal Fields Generator** to identify a meaningful CV term and to create the base field class, widget, and formatter.

#### Training a New Developer

A new developer has joined the team with some experience in bioinformatics, but no knowledge of Drupal, Chado, or Tripal.

The junior developer’s first task is to install a personal site using **TripalDock**. The site serves as a sandbox which can be easily destroyed and rebuilt, allowing them to learn and make mistakes without disrupting the deployed live site. It also allows them to begin learning Tripal quickly, without wrestling with system administration tasks (creating a virtual machine, webserver, etc). Next, they would practice loading data into their Tripal site using **DevSeed**. The fully documented dataset will teach them how Tripal imports different types of data, and how Chado stores it as well as how to navigate the administrative pages of a Tripal website.

Once their site is set up, the new developer is tasked with creating a simple module to learn the Tripal framework. They use the **Tripal Test Suite** setup command to give their project a functional PHPUnit testing and Travis CI environment on GitHub, without spending weeks understanding how to structure their tests, access their Drupal site within the tests, or create a Travis build with a functional Drupal. When they contribute code to other modules or to Tripal core, the test structure will match their learning module. Their learning module would use **Tripal Fields Generator** to build their first fields, automatically creating the necessary files in the correct location, with the required methods to fill out.

As the developer learns, they may load data into the development or production site as the wrong sub-type. Developers from a computer science background might be overwhelmed with the number of chado feature types to keep track of: mRNA, mRNA_contig, polypeptide, CDS, 5’ UTR, 3’ UTR, exon, intron, and gene. Tripal Alchemist makes correcting these mistakes trivial: any entities using the same base table can be easily converted, saving the team from deleting the offending entries and reloading.

When attempting to fix a problem on an existing Tripal module the team maintains, they introduce a crippling bug on the release branch. The continuous integration that the team set up with **Tripal Test Suite**’s notifies the team that the module no longer installs and tests do not pass, and the team is able to revert the change. They instruct the new member on using GitHub best practices (branches, pull requests) to contribute, and Continuous Integration tests on branches and pull requests allows the developer to submit functional code. Their own learning module used the same structure and framework for tests, so they will be able to write their own tests for their contribution.

## Conclusion

The release of Tripal 3 has unlocked new possibilities for community genomics websites. As with many projects, the limiting factor for a Tripal site is funding and, relatedly, personnel. With smaller development teams, tools that bridge skill set gaps, either in the biological or computer sciences, can greatly enhance productivity. We have introduced a suite of utilities that will facilitate working on Tripal 3 sites and developing Tripal 3 modules. These tools encourage best programming practices for module and site development, and ease the Tripal and Chado learning curves.

## Acknowledgements

We acknowledge the entire Tripal software developers community, particularly Stephen Ficklin (Washington State University) and Lacey-Anne Sanderson (University of Saskatchewan), for their leadership of the core Tripal infrastructure and the community of Tripal software developers. This work was supported by the National Science Foundation under grants #1443040 and #1444573.

